# 4D mitochondrial network assumes distinct and predictive phenotypes through human lung and intestinal epithelial development

**DOI:** 10.1101/2025.04.09.648043

**Authors:** Gillian McMahon, Dhruv Agarwal, Mehul Arora, Zichen Wang, Hiroyuki Hakozaki, Johannes Schöneberg

**Author notes:** Address correspondence to Johannes Schöneberg, Ph.D., Departments of Pharmacology and Chemistry and Biochemistry, University of California, San Diego, Center for Neural Circuits and Behavior, Room 106B, La Jolla, CA 92093.

## Abstract

Mitochondria form a dynamic three-dimensional network within the cell that exhibits a wide range of morphologies and behaviors. Depending on cell state, cell type, and cell fate, a cell’s mitochondrial phenotype might range from relatively isolated mitochondrial segments to complex branching networks, and from stationary mitochondria to highly motile structures. While isolated mitochondrial phenotypes have been described for a subset of cell states, types, and fates, an integrated map of how mitochondrial phenotypes change over the full course of tissue development has so far been lacking. Here, we identify the mitochondrial phenotypes that appear throughout the course of lung and intestinal epithelial development from stem cells to differentiated tissue. Using human stem cell-derived intestinal and branching lung organoids that mimic developing human organs as model systems, we extract and analyze key mitochondrial biophysical phenotypes in human development. To achieve this, we employ lattice light-sheet microscopy (LLSM), which enables high-resolution, 4D (x, y, z, time) imaging of mitochondria in organoid tissues with minimal damage to the sample. We image at key developmental time points from stem cell differentiation into mature organoid tissue. For data processing, we utilize the MitoGraph and MitoTNT software packages along with our developed custom computational tools. These tools allow for automated 4D organoid to single cell image processing and quantitative 4D single cell mitochondrial temporal network tracking. This work represents the first 4D high spatiotemporal-resolution quantification of live human organoid tissues at the single-cell level through development. We identified distinct mitochondrial phenotypes unique to each organoid type and found correlations between mitochondrial phenotypes, cellular age, and cell type. Furthermore, we demonstrate that mitochondrial network characteristics can predict both organoid type and cell age. Our findings reveal fundamental aspects of mitochondrial biology that were previously unobservable, offering new insights into cell-type-specific mitochondrial dynamics and enabling new findings in relevant human model systems. We believe that our findings and methods will be essential for advancing 4D cell biology, providing a powerful framework for characterizing organelles in organoid tissues.

## Introduction

Mitochondria have long been recognized as the primary energy source in cells, responsible for generating approximately 90% of the energy required for cellular function^1,2^. However, recent research has revealed that mitochondria play a far more complex role beyond energy production, contributing to diverse cellular processes such as apoptosis, immune signaling, and metabolic regulation^3–7^. Notably, mitochondria have been increasingly recognized for their roles in cell differentiation pathways^8^ where they not only undergo drastic changes in morphology and dynamics, but the success of assuming the required morphologies and dynamics of a new cell type controls if cellular differentiation proceeds^9^. As such, mitochondria have recently been recognized to be the gatekeepers of cellular differentiation in multiple cell types.

One particular instance where mitochondria have been shown to control cell fate decisions^7,10^ and where the morphological changes throughout differentiation are reportedly large^10^ is the differentiation of human stem cells into polarized epithelia. The two major epithelia of that class are the lung and intestinal epithelium. A previous study has even demonstrated the ability to switch cells committed to lung fate to an intestinal fate, suggesting a link between the two lineages^11^. The development of these tissues in humans is well characterized at the genetic, cellular, and tissue levels^12,13^. Early in human development, gastrulation leads to the formation of the three germ layers. The innermost layer, the endoderm, gives rise to many organs in the body, including those in the respiratory and digestive tracts such as the lungs and intestines. Cells in the anterior portion of the endoderm become the definitive endoderm (DE) and further differentiate into the foregut, midgut, and hindgut. The foregut pathway gives rise in part to the lungs and the mid/hindgut pathway gives rise in part to the intestines^14^.

There is a gap in knowledge as to the specific roles of mitochondria in differentiation pathways. It is widely accepted that generally as cells differentiate and become more specialized, they rely more on the mitochondria and oxidative phosphorylation (OXPHOS) for their source of energy^15^, shown by energetic assays that study oxygen consumption. It is also widely accepted that as cells become more specialized, their mitochondria become longer and more networked^16^. Beyond observing trends in development, researchers have explored how mitochondrial dysfunction affects specific steps in differentiation^8^. One example is that inhibition of mitochondrial restructuring in the intestinal epithelium has been shown to disrupt the differentiation of intestinal stem cells into specialized cell types, such as Paneth cells^10,17^.

Currently, there are three limitations that prevent us from fully understanding the role of mitochondria in development: 1) the traditional reliance on optical sections, i.e. two dimensions (2D), to understand the four dimensional (4D, x,y,z,time) character of mitochondrial phenotypes, 2) the study of human cells outside their three dimensional (3D) tissue context, i.e. as adherent cultures rather than in a tissue, and 3) the study of limited differentiation snapshots, i.e. differentiation from one cell type to the next cell type, without taking the entire process of differentiation from stem cells to mature tissue into account.

Optical sectioning can leave out valuable information relating to the 4D mitochondrial network^18^. Thus, understanding 3D mitochondrial network dynamics is vital to understanding human development, how development can be disrupted, and creating new therapeutic strategies. The recent development of 4D mitochondrial tracking tools by us (MitoTNT^19^) and others (e.g. Mitometer^20^, Nellie^21^) allows the capture of the 4D nature of mitochondria for the first time. In this study, we aim to consistently use 4D mitochondrial tracking to quantify the role of mitochondria in development.

The development of human stem cell-derived organoid systems have enabled researchers to model human tissue formation and function in a controlled, three-dimensional environment more similar to human organs than conventional monolayer systems^14,22^. Up until recently, such organoid systems could largely only be studied using static endpoint techniques, such as RNA sequencing and immunofluorescence imaging of fixed and cleared samples^23^. While these approaches provide valuable insights into gene expression and protein localization, they fail to capture the dynamic processes occurring at the subcellular level, such as mitochondrial dynamics. Traditionally, confocal microscopy has been the primary method for live imaging of organoids^24^. This technique achieves high resolution imaging by illuminating the entire sample and rejecting out-of-focus light to generate volumetric datasets, but is inherently limited by phototoxicity and photobleaching that restricts the duration and frequency of imaging sessions^25^. As a result, dynamic processes, such as mitochondrial remodeling during differentiation, remain difficult to capture beyond a few volumetric frames.

The recent development of lattice light-sheet microscopy (LLSM) offers a transformative solution to these challenges^26^. When combined with adaptive optics^27^, this advanced imaging technique provides high-resolution low-phototoxicity volumetric imaging of organoid tissues at high temporal resolution for extended periods of time^28^.

In this study, we use LLSM to quantitatively analyze the 4D mitochondrial network through lung and intestinal development. Specifically, we utilized human induced pluripotent stem cells (hiPSCs) expressing endogenously tagged mitochondria and cell membrane fluorescent markers to visualize mitochondrial dynamics throughout the development of hiPSCs into intestinal and lung epithelial organoids, 90 and 150 days later respectively. We generated and analyzed more than 70 4D movies, capturing single cells in organoids with the temporal resolution necessary to study dynamic mitochondria information. Our approach allowed us to segment individual cells, track their mitochondrial activity over time, and perform detailed 4D analyses of human organoid tissue at the single-cell level.

Our findings reveal that mitochondria shift from interconnected networks in hiPSCs to more linear and motile structures in organoids, with intestinal organoids displaying an intermediate phenotype between hiPSCs and lung organoids. Furthermore, we demonstrate that mitochondrial features can serve as predictive biomarkers of cell type and maturation. This study establishes a new framework for understanding mitochondrial dynamics in human development and highlights their potential as indicators of cellular identity and function.

## Results

### 4D Single Cell Live Organelle Quantification in Human Organoids is Enabled by Custom Lattice Light Sheet Microscopy and Custom Software

In order to study 4D mitochondrial phenotypes through epithelial development we used established protocols (see Methods) to differentiate hiPSCs into branching lung organoids (BLOs) and intestinal organoids (IOs), thereby recapitulating human development (Fig. 1a-d). Briefly, hiPSCs (Fig. 1e,j) were differentiated into definitive endoderm (DE), where we saw the first 3D structures starting to form (Fig. 1f,k). As expected, buds arose from these 3D structures and started floating in the media (Fig. 1g,l). BLOs and IOs at this stage appeared similar in structure, with buds of single epithelial cell layers starting to form and expand. For BLOs, these buds were moved to suspension culture where the epithelial branches continued to develop (Fig. 1h). These more branched buds were then embedded in Matrigel for maturation (Fig. 1i). For IOs, the buds were immediately moved to a Matrigel dome, where they grew more spherically (Fig. 1m). IOs are made up of intestinal stem cells (CDX2+) (Fig. 1n), which are more proliferative than lung progenitor cells (NKX2-1+) (Fig. 1l), thus IOs expanded to fill the Matrigel dome in 7-10 days and required passaging (Fig. 1n).

**Fig. 1:**
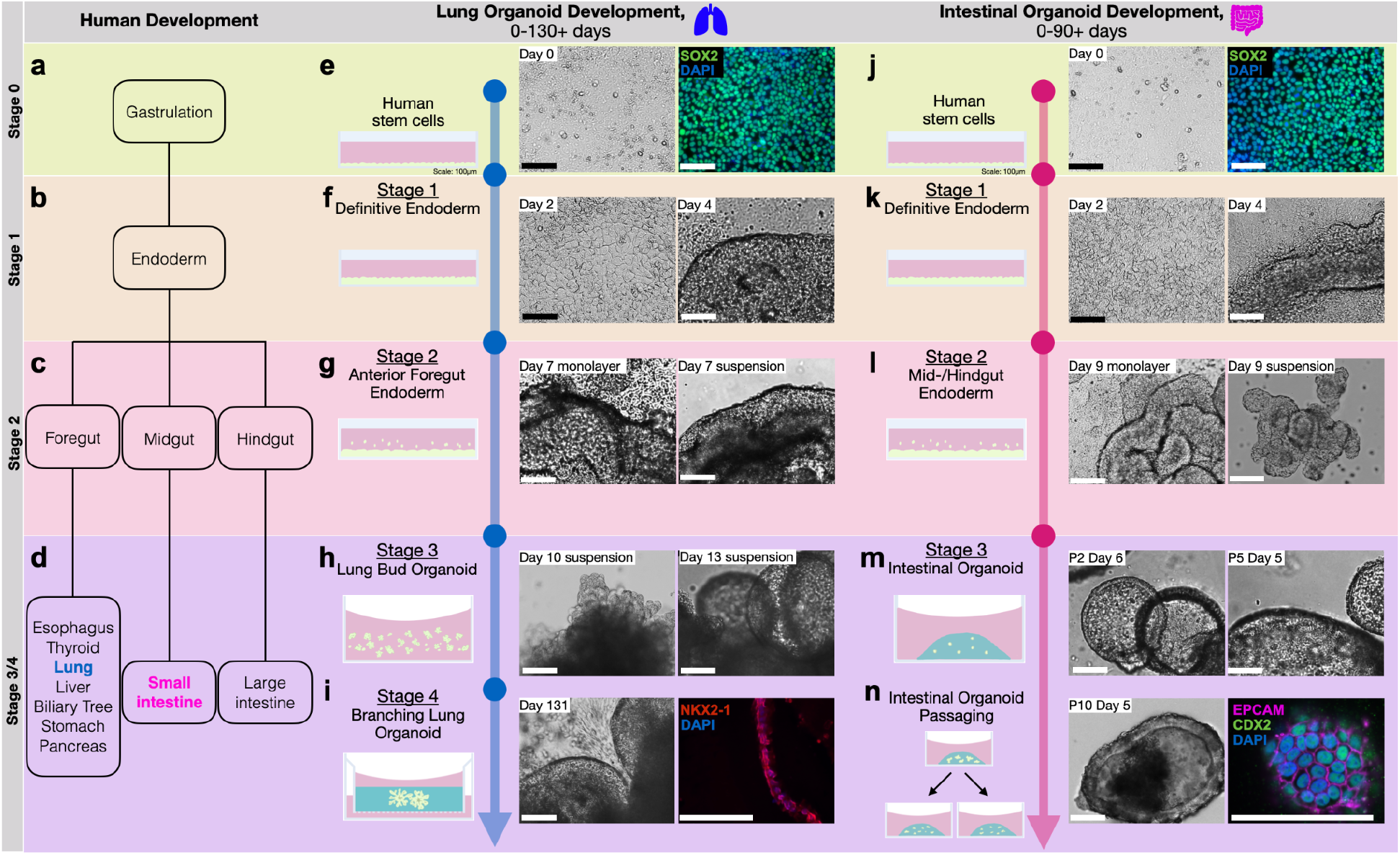
Human lung and intestinal organoid differentiation recapitulates human developmental stages. **a-d**, Most human organs result from the endoderm differentiation pathway, notably the lung from the anterior foregut and the intestines from the mid/hindgut. **e-i**, hiPSCs can be differentiated into mature BLOs that mimic epithelial lung progenitor cells (NKX2-1+). **j-n**, hiPSCs can be differentiated into IOs that mimic the regenerating intestinal epithelium (CDX2+). *Scale: 100µm*.

The differentiation protocols of BLOs and IOs are similar in the endoderm stages of development (Fig. 1f,k and 1g,l) and their single cell epithelial layer structure (Fig. 1i,n), however one key difference is their proliferation rate. In maturation, the BLOs that were initially embedded in Matrigel continued growth while the IOs that were initially embedded in Matrigel were split up every 7-10 days. Thus, mature BLOs were largely composed of the same cells that were initially embedded and mature IOs were largely composed of new cells originating from fragments of IOs after passaging. Consequently, IOs exhibit more “stemness” through their proliferation rates.

In order to study the 4D mitochondrial phenotype at every stage of BLO and IO organoid development, the following steps were performed as a pipeline (Fig. 2). First, hiPSCs, tagged with Tom20-mEGFP (mitochondria) and CAAX-TagRFP (cell membrane), were cultured (Fig. 2a) and differentiated into BLOs and IOs using established protocols (Fig. 2b). At specific developmental timepoints, the organoids were live imaged using LLSM (Fig. 2c) and the raw 4D data pre-processed using LiveLattice^29^ (Fig. 2d).

**Fig. 2:**
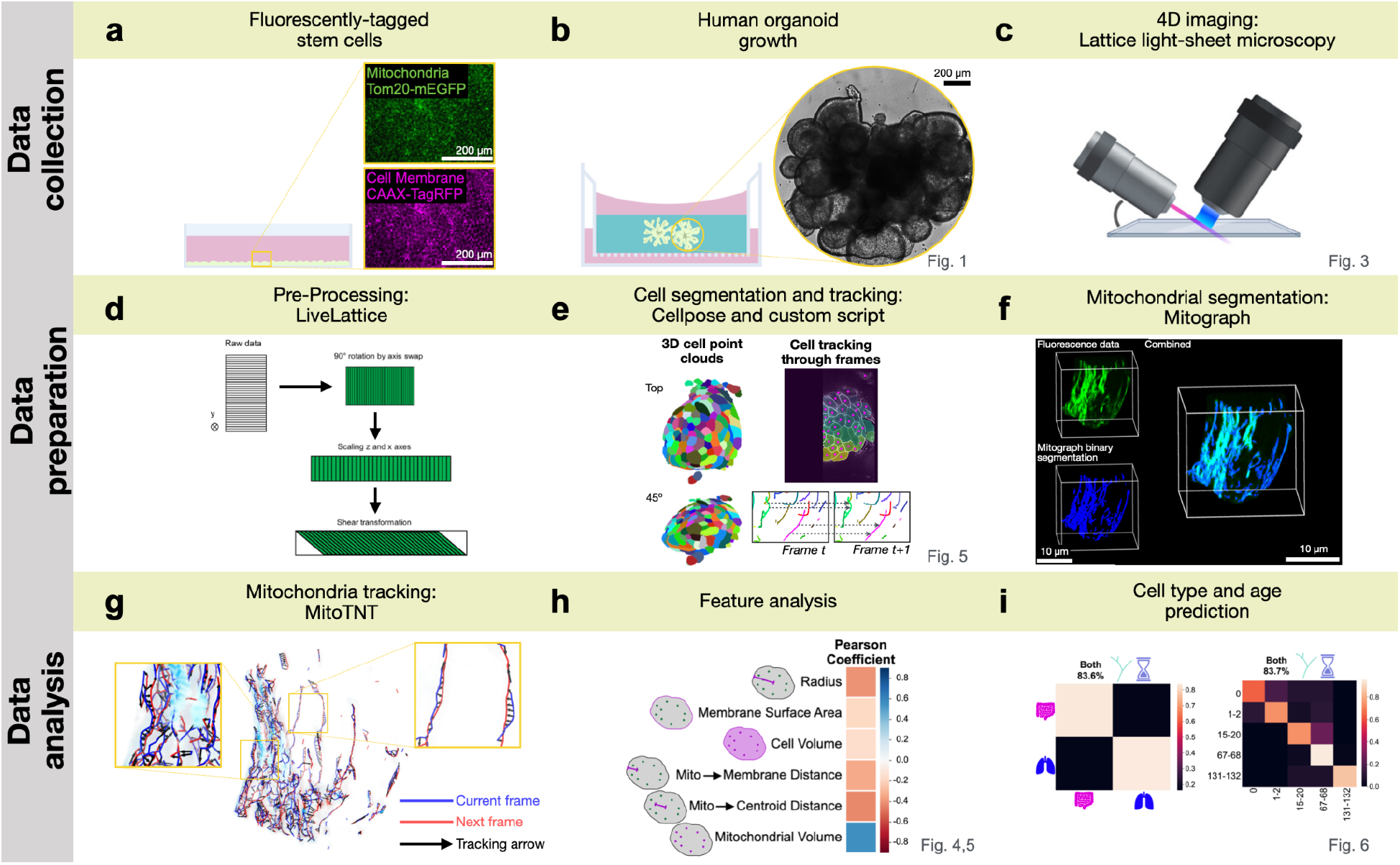
A custom 4D single cell organoid tissue data processing pipeline enables organelle quantification in single cells that are part of the organoid tissue. **a**, hiPSCs with the mitochondria and cell membrane fluorescently tagged were grown into **b**, human organoids. **c**, At different developmental timepoints, cells and organoids were 4D imaged using LLSM. **d**, Resulting data was pre-processed using lab-developed LiveLattice methods. **e**, Single cells were segmented in 3D and tracked across frames to extract 4D single cell data. The mitochondrial network was **f**, segmented and **g**, tracked to quantify fluorescent data. Mitochondrial network and single cell features were **h**, analyzed and **i**, used to predict organoid type and lung cell age.

In order to assign the 4D mitochondrial phenotypes to individual cells, the organoid tissue was segmented using the cell membrane channel (Fig. 2e). Briefly, every 3D frame of the 4D movie was segmented using Cellpose^30^ and the centers of these cells were tracked over time as 3D point clouds using DBSCAN^31^. The results are segmented 4D movies of single cells in the organoid tissue.

The 4D mitochondrial phenotype was extracted from each of these single organoid cells as follows: MitoGraph^32^ was used to segment the mitochondria using the TOM20-mEGFP fluorescence signal and create a mitochondrial skeleton (Fig. 2f). This skeleton was tracked across frames with MitoTNT (Fig. 2g). The result is a segmented and tracked 4D mitochondrial network for single cells in the organoid tissue.

To compare mitochondrial morphologies and motilities in different organoid types and in different developmental time points, quantitative mitochondrial features were extracted (Fig. 2h). In the last step, machine learning tools were applied to the extracted features to compare organoid type and cell age against each other. (Fig. 2i).

We were able to study subcellular dynamic information in single cells in live human organoids and used this information to predict cell type and age.

### Mitochondrial Networks Change Primarily According to Cell Age and Secondarily According to Cell Type

Using the described pipeline, we imaged the 4D mitochondrial network during the differentiation from hiPSCs through DE into BLO with LLSM at key developmental stages (Fig. 3a). Using the cell membrane, single cells could be extracted from each dataset to show the mitochondrial network dynamics from individual cells in the monolayer or BLO tissue environment (Fig. 3b). The same was done for IOs: we imaged differentiation from hiPSCs through DE into IOs (Fig. 3c). Single cells were extracted for these datasets as well (Fig. 3d). In comparing full datasets, we observe the shift from monolayer cells to epithelial layers of organoid tissues. The lumen within the epithelial layer in lung d68 and d131 and in intestine d49 is clearly visible. We see more compact packing of the cells in organoid tissue.

**Fig. 3:**
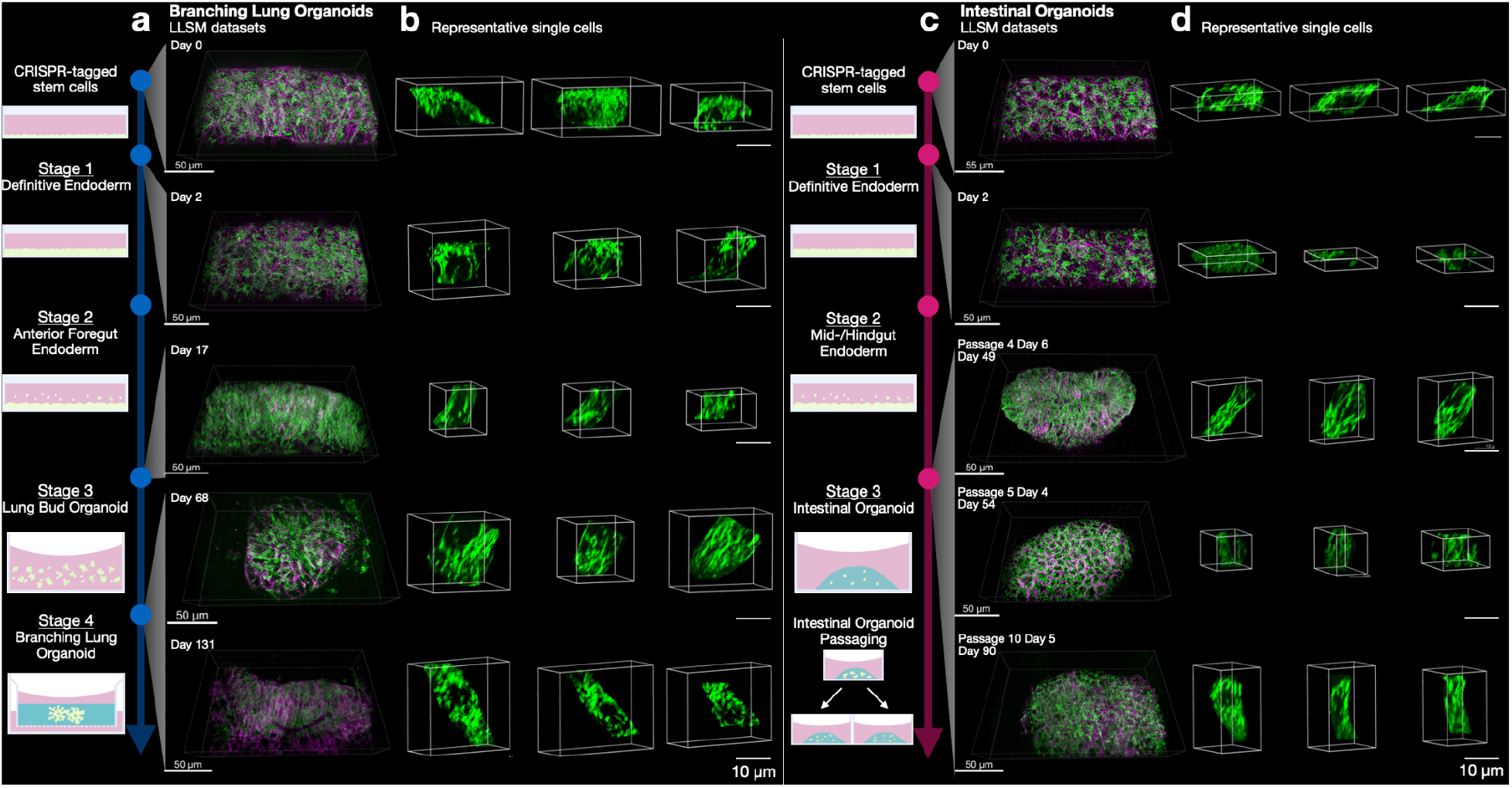
4D Lattice light-sheet imaging of mitochondria in lung and intestinal organoids results in large imaged tissue sections that can be segmented into single cell 4D datasets. **a**, Key developmental time points in the differentiation of hiPSCs into BLOs were imaged with LLSM. **b**, Single cells were segmented from each dataset, revealing the morphological transition from stem cells to mature lung cells. **c**, Key developmental time points in the differentiation of hiPSCs into IOs were imaged with LLSM. **d**, Multiple single cells were segmented from each dataset, revealing the morphological transition from stem cells to intestinal cells.

In lung development, we observe wide early development-stage cells (hiPSC d0, DE d2), first becoming smaller and more compact (anterior foregut endoderm d17), and eventually assuming long and thin cell morphologies. Mitochondrial networks are at first distributed throughout the cytoplasm (d0, d2) and move closer to the cell membrane as development progresses (Fig. 3b,d).

To quantify the 4D mitochondrial network across organ types and developmental stages, mitochondrial network features were extracted from 4D LLSM data. Briefly, in an ROI containing multiple cells, mitochondria were segmented and skeletonized (Fig. 4a). For each frame these skeletons were quantified to study i) the total length of all mitochondrial network fragments, and ii) the branch lengths within these fragments, i.e. the distance between ends and/or junctions within the network fragment (Fig. 4b). To quantify on average how far the mitochondrial network moves over time, we studied the mean square displacement (MSD) of network fragments and branches using MitoTNT (Fig. 4c). The MSD quantification demonstrates on average how far the branch moves during the given frame interval. To understand how each metric changed with age, we calculated the Pearson correlation coefficient and the slope of the regression fit for metric averages for all ROIs from each age (Fig. 4d).

**Fig. 4:**
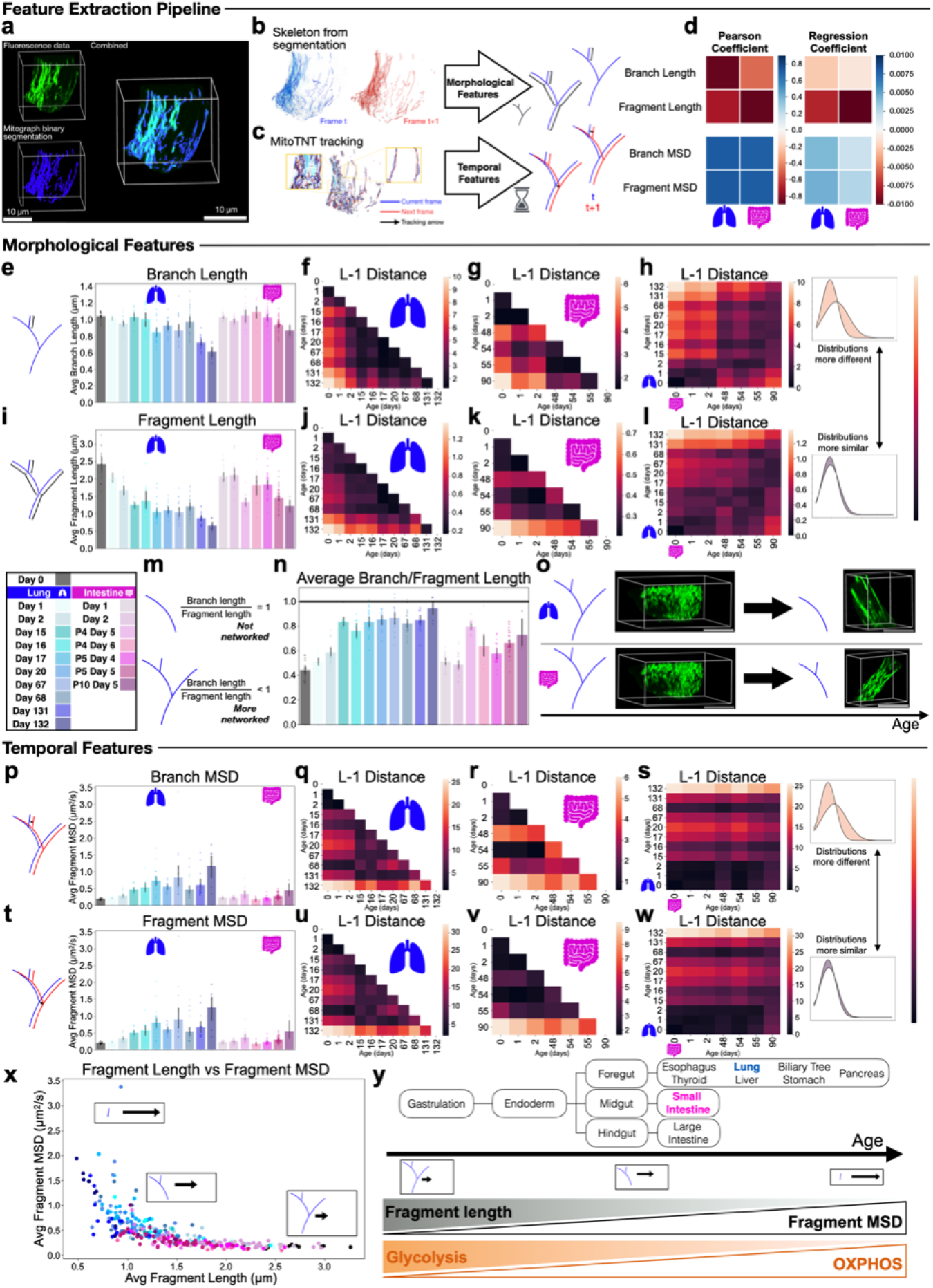
Global quantification of mitochondrial dynamics show age-related and cell type-specific trends. **a**, Fluorescent mitochondrial data was segmented with MitoGraph to generate skeletons of the mitochondrial network for each frame. Skeletons were analyzed with MitoTNT to study **b**, morphological and **c**, temporal features of the network for each condition. **d**, Morphological and temporal features are correlated with age. **e**, Mitochondrial branch length is relatively consistent, until days 131-132 of BLO growth. **f**,**g**, Mitochondrial branch length distribution in days 0-2 of development differs the most from other developmental timepoints. **h**, The distribution of mitochondrial branch length values is most different when comparing young and old cells and stays more similar when comparing different organoids across age. **i**, Mitochondrial fragment length decreases when lung and intestinal organoids mature, with fragments in BLOs decreasing more. **j**,**k**, Mitochondrial fragment length distribution in the most mature cells differs the most from other developmental timepoints. **l**, The distribution of mitochondrial fragment length values is most different when comparing to mature lung cells and stays more similar when comparing different organoids across age. **m**, A branch length to fragment length ratio =1 represents no networking, while a ratio <1 represents some networking. **n**,**o** Ratios of average branch length to average fragment length show levels of mitochondrial networking. *Scale: 10µm* **p**, Mitochondrial branch MSD increases as BLOs mature. **q**,**r**, Mitochondrial branch MSD distribution in the most mature cells differs the most from other developmental timepoints. **s**, Organoid type differentiates branch MSD distributions more than age. **t**, Mitochondrial fragment MSD increases as BLOs mature. **u**,**v**, Mitochondrial fragment MSD distribution in the most mature cells differs the most from other developmental timepoints. **w**, Organoid type differentiates fragment MSD distributions more than age. **x**, Shorter fragments move more in a given amount of time than longer fragments. **y**, Mitochondrial fragments get smaller and move more as cells mature.

We observed high linear correlation with age for all metrics in both lung and intestine organoids, where mitochondrial network lengths decreased with age (Fig. 4d top left) and mitochondrial motilities increased with age (Fig. 4d bottom left). The fragment length decreased more with age compared to branch length in both organoid systems (Fig. 4d top right). Branch and fragment MSD increased with age more in the lung than the intestine (Fig. 4d bottom right). Overall, we saw that mitochondrial metrics increase and decrease linearly with age in both lung and intestinal organoids. Fragment length changes more with age than branch length. Mitochondria in BLOs move more with age than mitochondria in IOs.

To further explore these trends, we studied the relationship between distributions of these values and organoid type and age.

First, we focused on mitochondrial branch lengths, the fundamental building blocks out of which mitochondrial networks are constructed. We saw a slight decrease in branch length in lung and intestine mitochondria, but branches in BLOs get shorter than branches in IOs (Fig. 4e). Branch lengths in IOs decrease 1.2-fold ±0.1, branch lengths in BLOs decrease 1.7-fold ±0.3. To study distributions more closely, we compared the L-1 distance for metric histograms between every sample, with lighter boxes representing greater difference and darker boxes representing less difference between two distributions. We first compared the branch length values between every age in BLOs (Fig. 4f). We observed that the most drastic difference is between hiPSC/DE cells (d0-2) and more mature cell types (d15-132) in the lung. Mitochondria in the more mature cell types have more similar distributions of branch lengths to each other. Similar trends were seen in IOs (Fig. 4g): hiPSCs and DE (d0-2) cells’ branch length distributions are similar to each other, but different from mature IO (d48-90) cells. We also compared branch length distributions between organoid types through age and saw that hiPSC/DE, young organoids, and mature organoids have distributions that are more similar regardless of organoid type, shown by the darker squares on the diagonal (Fig. 4h). We also observed that young intestine cells are more different from old lung cells than young lung cells are from old intestine cells.

Next, we focused on mitochondrial fragment lengths, the overall length of the mitochondrial networks built from the branches discussed in the previous section. Where we had already seen a decrease in branch length before, we now saw an even more pronounced decrease in fragment length through age in both organoid types (Fig. 4i). As with branch length, mitochondria in BLOs have shorter fragment lengths than in IOs, where BLO fragment lengths decrease by 3.7-fold ±1.0 in length through development as compared to 2.0-fold ±0.5 in IOs. In comparing fragment length distributions between lung ages, we saw the most different distribution is between the oldest organoids and the rest of the data (light rows in Fig. 4j). In IOs, again the oldest organoids are the most different from the rest of the data (light rows in Fig. 4k). When comparing the two organoid systems’ fragment lengths, we found that similar relative ages show similar fragment length distributions (Fig. 4l). Additionally, the oldest BLOs exhibited the most distinct fragment length distributions compared to other samples, shown by the brightest row in the heatmap at lung d131/d132.

To investigate the extent to which a mitochondrial fragment is networked, we next calculated the ratio of branch length to fragment length (Fig. 4m). A ratio of 1 signifies that the average branch length and average fragment length are equal, meaning the fragment lacks additional branches. A ratio below 1 suggests that the average branch length is shorter than the average fragment length, indicating a higher degree of branching and networking within the fragment.

We found that stem cells had the lowest ratio, indicating the most networked mitochondrial fragments (Fig. 4n, grey bar). As cells differentiated into BLOs, the ratio increased and approached one (Fig. 4n, blue bars). As cells differentiated into IOs, the ratio increased, but not as much as in the lung samples (Fig. 4n, magenta bars). Overall, as cells mature, their mitochondria were observed to get less networked. In fact, we observe mitochondria in lung organoids to approach no networking, instead creating strands of long mitochondrial filaments near the membrane along the major axis of the cell (Fig. 4o).

After static morphological features showed clear differences in mitochondrial phenotypes between organoid types and across tissue maturation, we hypothesized that similar differences could be observed in the temporal mitochondrial features. LLSM allows us to image live cells and organoids with high temporal resolution to track individual mitochondria branches and quantify their movement. We tracked mitochondrial motility at a volumetric frame rate of 10.7 seconds/frame and quantified it.

We observed that the motility of mitochondrial branches increases with age by 6.2-fold ±2.8 in lung and 2.4-fold ±1.1 in intestinal organoids (Fig. 4p). The distribution of branch MSDs in the lung is most different when comparing organoids to early cells, with the greatest difference being between d0,1,2 and d132 (three white squares in Fig. 4q). Additionally, distributions between d132 and d0,1,2 (Fig. 4p, first three columns) are distinct from the rest of the data. Branch MSD distributions in IOs are distinct from each other, except within the first few days and d54 (Fig. 4r). Comparing mitochondrial motility in the two organoid systems (Fig. 4s), the differences in distributions are higher between organoids than across ages, indicated by the distinct horizontal lines showing that mitochondria in BLOs have different distributions from IOs regardless of intestinal organoid age. Notably, lung d0,1,2 have branch MSD distributions that are much more similar to all ages of intestinal organoids compared to all other lung ages.

When moving from branch motility to fragment motility, we observed very similar trends (Fig. 4t). Mitochondrial fragments move more as hiPSCs differentiate into BLOs and IOs (6.3-fold increase ±2.8 in lung, 2.8-fold increase ±1.6 in intestine). Comparing the distributions of fragment MSDs in lung shows that the oldest BLOs have the most distinct distribution, and d0,1,2 have distributions different from other ages (Fig. 4u). Fragment MSDs in the intestine have the most distinct distribution at the most mature stage (Fig. 4v). These d90 organoids are also more different from other ages in terms of fragment MSD than they are in branch MSD. We see this when we compare the scale of the heatmap in Fig. 4v vs 4r, the rest of the heatmap is similar in value and the d90 row is much higher. As with branch MSD, fragment MSD across organoid types differs primarily according to organoid type and secondarily according to age (Fig. 4w). There is little difference in distributions between the d0,1,2 lung cells and all IOs, indicating similarity in mitochondrial motility between hiPSCs/lung DE and IOs.

To investigate the relationship between the size of a mitochondrial fragment and its motility, we compared the average fragment length to the average fragment MSD. We observed an inverse correlation: smaller fragments showed higher MSDs, and larger fragments showed lower MSDs (Fig. 4x). This is consistent with first principles, as larger structures require greater effort to traverse a given distance compared to smaller ones. When we consider ages, we see a continuum right to left along the x-axis of hiPSCs into young, then mature IOs, into mature BLOs.

In summary, we observe that hiPSCs have the longest, most connected network that moves the least and BLOs have the least connected network that moves the most, with IOs residing in the middle (Fig. 4y). Since proliferative potential is also the highest in hiPSCs and the lowest in BLOs with IOs residing in the middle, and the metabolic state of these cells is most shifted to glycolysis in hiPSCs and most to OXPHOS in BLOs with IOs residing in the middle ^33,34^, our data suggests there is not only a physical relationship between a fragment of mitochondria and how it moves, but also that the proliferative potential and cellular metabolic demands can influence the form and motility of its mitochondrial network (see Discussion).

After observing that the mitochondrial network assumes morphologies and display motilities that form distinct signatures across development, we next hypothesized how these mitochondrial network morphologies and motilities are related to the cells in which they are residing. We segmented our 4D LLSM organoid data (Fig. 5a) into individual cells using Cellpose and DBSCAN, (Fig. 5b) which were then tracked over time using cell centroids (Fig. 5c). The final result was a 4D mask of the volume of each individual cell (Fig. 5d) in which mitochondria properties could now be quantified in relation to the surrounding cell (Fig. 5e).

**Fig. 5:**
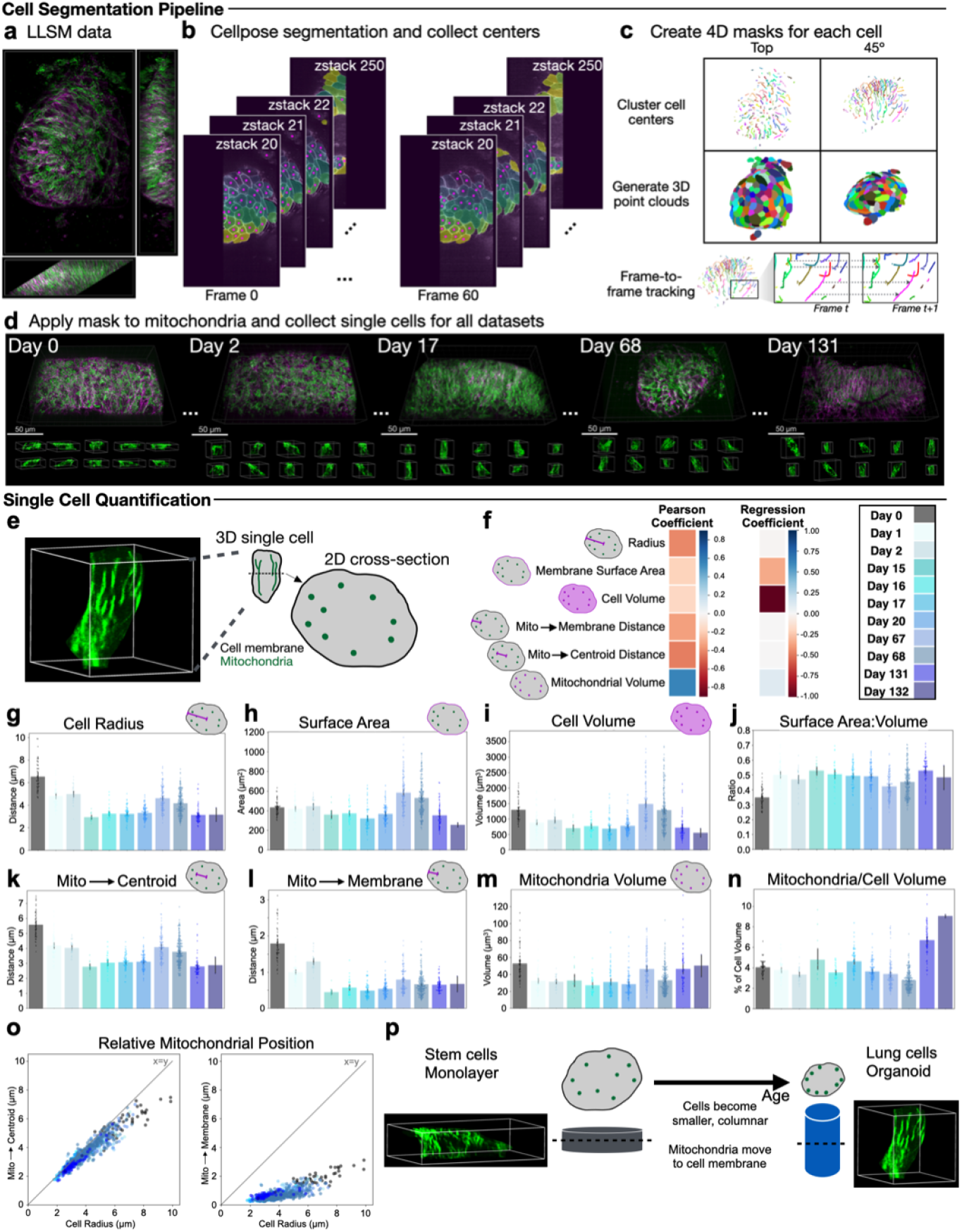
Single cell quantification of mitochondrial dynamics and cellular features show trends in lung epithelial cell development. **a**, Single cells from 4D fluorescent data of mitochondria and cell membrane can be segmented to quantify organoid tissue at single-cell resolution. **b**, The cell membrane was segmented to create cell masks and extract cell centroids. **c**, Centers were clustered using DBSCAN to connect masks in 3D space and tracked across consecutive frames to generate 4D cell segmentations. **d**, Cell masks were generated for cells in each dataset, which were applied to mitochondria data, resulting in numerous single cell movies of the mitochondrial network from live human organoid tissue. **e**,**f**, Single cell-resolution data enables analysis of cell morphology and mitochondrial relationships within individual cells. **g**, Cell radius decreases as lung cells mature. **h**, Cell surface area slightly decreases with age, but increases in d67/68 cells. **i**, Cell volume has similar trends to cell surface area, with greater volume in hiPSCs. **j**, The SA:V ratio of the cell is lowest in hiPSCs, then stays consistent. **k**, The mitochondria are closer to the centroid as cells mature, on average. **l**, The mitochondria are closer to the membrane in BLOs, on average. **m**, The volume of mitochondria in cells stays consistent. **n**, The mitochondrial volume:cell volume ratio increases in the oldest lung cells. **o**, Mitochondria in stem cells are more distributed around the cell, while mitochondria in lung cells hug the membrane. **p**, On average, cells start flat with a well-distributed mitochondrial network and mature into tall and skinny lung cells with mitochondria that hug the membrane.

We focused on three cell-centric properties (cell radius, cell surface area, and cell volume) and three mitochondria properties in relation to the cell (mitochondria distance to the cell centroid, mitochondria distance to the membrane, and mitochondrial volume) to quantify the relation between cell and mitochondria through the development of BLOs.

We found that all studied metrics decreased with developmental age except mitochondrial volume (Fig. 5f blue box). Cell surface area and cell volume had the lowest linear correlation and the greatest decrease with age compared to other metrics, with cell volume decreasing the most (Fig. 5f second and third rows).

Focusing on cell-centric metrics, we observed through BLO development that the cell radius was largest in hiPSCs, decreased in DE, and decreased even more in organoids (Fig. 5g). Cell surface area decreased with age, with the exception of d67,68 cells (Fig. 5h). These cells were observed to be generally larger, as seen in Fig. 5d. Cell volume followed a similar trend, however hiPSCs were observed to be greater in volume in comparison to the rest of the data (Fig. 5i). This could be better observed in the surface area:volume (SA:V) ratio (Fig. 5j). We saw an increase in SA:V from hiPSC to DE and another increase from DE to BLO. D67,68 cells had a lower SA:V ratio than other lung cells.

Shifting our focus on mitochondria-centric metrics, we observed that the average distance of mitochondria to the centroid of the cell (for each z) decreased from hiPSC to DE to BLO by about half from 5.6±0.9µm to 2.8±0.7µm (Fig. 5k). These relationships were the same as those seen in cell radius in 5g. The average distance from mitochondria to cell membrane decreased from hiPSC to DE to BLO, where the average distance between mitochondria and membrane was less than 1 µm (Fig. 5l). The mitochondrial volume was largely consistent across ages (Fig. 5m). However, normalizing mitochondrial volume by cell volume shows proportionately more mitochondria in the most mature organoids (Fig. 5n).

Relating cell and mitochondria-centric metrics, we observed that the distance from mitochondria to the centroid increases with the cell radius, almost in a linear fashion (Fig. 5o left). In contrast, the distance from mitochondria to the cell membrane showed minimal increase with increasing cell radius (Fig. 5o right). Thus, mitochondrial distance to centroid is correlated with cell radius, while mitochondrial distance to membrane is correlated with cell type (Fig. 5o). Overall, we observed a trend of short and wide hiPSCs differentiating into tall and narrow lung epithelial cells in which an originally distributed mitochondrial network starts to shift closer and closer to the cell membrane with development (Fig. 5p).

### Cell Type and Age Can Be Predicted with High Accuracy Using Cell Morphology and Mitochondrial Morphology and Motility

After having observed clear trends in mitochondrial morphology, mitochondrial motility, and their relations to cell shape through epithelial development, we hypothesized that the dataset we collected would contain enough information to potentially predict cell type or organoid age. After compiling all our collected metrics (Fig. 6a), each metric was standardized through normalizing to its maximum value (Fig. 6b) and calculating averages and standard deviations, which we combined with the histogram features to construct a feature matrix for further analysis (Fig. 6c). With the feature matrix of data from lung and intestine ROIs, we first conducted principal component analysis (PCA) to determine the greatest sources of variation in the data (Fig. 6d). The first principal component (PC), which explains 25.4% of the variance, correlates with cell type (x-axis in Fig. 6d). From left to right on the x-axis, we move from hiPSCs to DE to intestine to lung.

**Figure 6:**
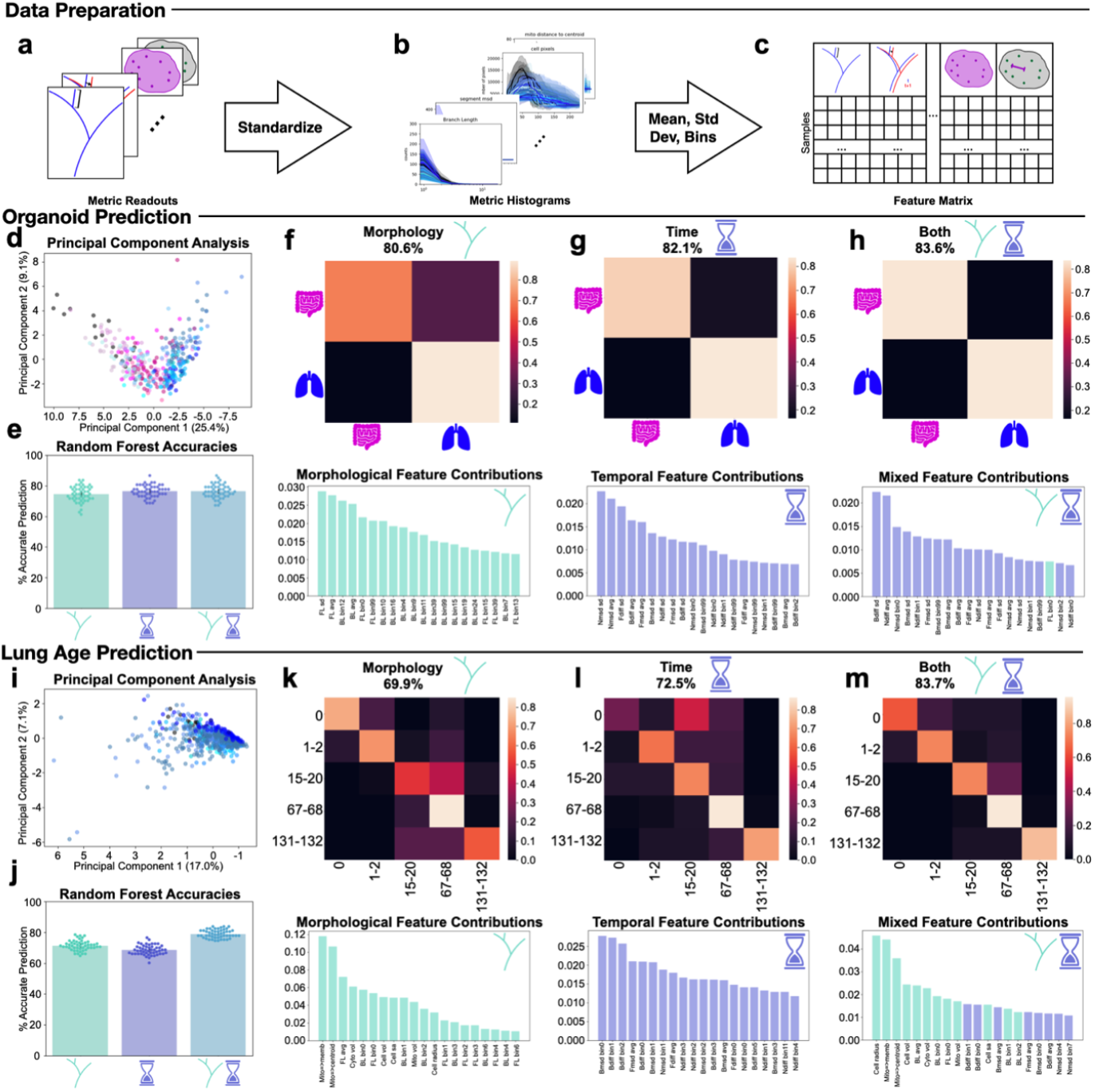
Organoid type and lung cell age can be predicted using mitochondrial network morphology and dynamics. **a**, Each quantification metric readout is stored in a distinct data structure and subsequently compiled for standardization. **b**, Histograms enable comparison across ages and standardize evaluation. **c**, Means, standard deviations, and distribution histograms for each metric make up a feature matrix to compare all features for all samples. **d**, PCA shows the greatest variation in data comparing organoids is the cell type the mitochondria belongs to. **e**, A random forest model was used to predict organoid type on 50 different train-test data splits with morphological, temporal, and morphological+temporal features. **f**, In an example run, morphological features yielded 80.6% accuracy in predicting organoid type. **g**, Temporal features yielded 82.1% prediction accuracy. **h**, Using both morphological and temporal features yielded 83.6% accuracy, with temporal features being the most important in decisions. **i**, PCA is used to separate individual lung cell ages. **j**, Using both morphological and temporal features in random forest prediction gives the highest accuracy. **k**, In an example run, morphological features yielded 69.9% accuracy in predicting lung cell approximate age. **l**, Temporal features yielded 72.5% prediction accuracy. **m**, Using both morphological and temporal features yielded 83.7% accuracy, with morphological features being the most important in decisions.

Seeing that our data has underlying variation across cell types, we attempted to predict organoid type with a random forest model by training it on 80% of the data and predicting on the unseen 20%. We ran the experiment on 50 different data splits using only morphological data, only temporal data, and both combined to ensure robust results across varying random forest seeds (Fig. 6e).

With only morphology, we predict organoid type with 80.6% accuracy (Fig. 6f). The most important features were mitochondrial fragment length standard deviation (SD), mitochondrial fragment length mean, and the mitochondrial branch length distribution.

With only temporal features, we predict organoid type with 82.1% accuracy (Fig. 6g). The most important features were node MSD SD and mean, and fragment diffusivity SD and mean.

With both morphological and temporal features, we predict organoid type with 83.6% accuracy (Fig. 6h). All except one of the top twenty most important features were temporal features, with the 18th most important feature being the mitochondrial fragment length distribution. The top features were branch diffusivity SD and node diffusivity mean. Since the accuracy was higher in Fig. 6g than 6f and the top 20 feature contributions were mostly temporal, we see that organoid type prediction relies heavily on temporal data.

Moving from ROIs to single cell BLO data, we first conducted PCA (Fig. 6i) and observed correlation between the first two PCs and cell age. Next, we attempted to predict the approximate cell age with a random forest model using only morphological data, only temporal data, and both morphological and temporal data on 50 different training and testing splits (Fig. 6j).

Using both temporal and morphological data yielded the highest accuracy, while the loss of morphological data yielded the lowest. With only morphology, we predicted approximate age with 69.9% accuracy (Fig. 6k). The most important features were average distance from mitochondria to membrane, average distance from mitochondria to centroid, and fragment length. With only temporal features, we predict approximate age with 72.5% accuracy (Fig. 6l) with the most important features being branch MSD and diffusivity distribution. With both morphological and temporal features, we predicted organoid type with 83.7% accuracy (Fig. 6m). The most important features were cell radius, average distance of mitochondria to cell membrane, and average distance from mitochondria to centroid.

Overall, we see that it is possible to predict organoid type and cell age with high accuracy. To succeed, both temporal and morphological data is required.

## Discussion

Here we present the first study of mitochondrial morphology and motility through the development from hiPSCs to human lung and intestinal organoids using LLSM. By leveraging 4D imaging and custom software, we provide insights into mitochondrial behavior during organoid development and differentiation and explore fundamental behavior of the mitochondrial network. We have three main findings: 1) mitochondrial network fragments decrease in length and increase in motility as cells differentiate, 2) mitochondrial characteristics of IOs are between those of hiPSCs and BLOs, and 3) the mitochondrial network can be used as a predictive biomarker of cell type and cell maturity.

Our first finding is that mitochondrial fragments decrease in length and increase in motility with organoid maturation.

Focusing on morphology, we observed a decrease in mitochondrial fragment length (Fig. 4d,4i) along with a decrease in mitochondrial networking (Fig. 4n) for both organoid types. In the literature, it has been reported that mitochondria actually increase in length from short, granular fragments in stem cells to long, tubular strands in specialized cells^16,35,36^. The current hypothesis is that, as cells differentiate from stem cells to specialized cells, they require more energy in the form of ATP and that this increased ATP demand can be better supplied by longer and more networked mitochondria^37^. There are two possible reconciliations for the reported findings and our data. First, not all specialized cells demonstrate this behavior of mitochondrial elongation^38^, so our finding might be dependent on our choice of model system of BLOs and IOs. Second, the majority of reports on mitochondrial morphology are based on 2D imaging sections of the 3D mitochondrial network. It has been reported that valuable morphological information can be gained when the mitochondrial network is imaged in 3D^18^. This 3D imaging is often done with electron microscopy, which provides detailed spatial information but by nature cannot be done on living cells^39–41^. Now, through the recent advent of LLSM, a new source of 3D mitochondrial information is available. We hope that a larger adoption of LLSM and similar technologies can shed more light on 3D mitochondrial network morphology.

Focusing on motility, we observed an increase as cells became more specialized for both tested organoid types (Fig. 4p,4t). This is consistent with observations that increasingly specialized cells rely more on OXPHOS^15^ and that in such cells, mitochondria are transported to sites where ATP demand is high^42–44^. In contrast, in stem cells, where glycolysis is dominant^15^, the location of the less used mitochondria are therefore less controlled and less motile.

In fact, our increasing motility with increasing specialization observation is consistent with a model where increasing specialization simultaneously leads to a decrease in mitochondrial fragment length, as observed in our data as well. Smaller mitochondrial fragments are easier to move along the microtubule cytoskeleton. Indeed, a size limit has been reported for kinesin- and dynein-mediated transport of mitochondria^45^. Our results, for both organoid models, support this model.

Our next finding is that the mitochondrial network in IOs has phenotypes that lie between those of stem cells and BLOs. In the human body the intestinal epithelium is continually proliferating, and in adults is the most rapidly proliferating organ^46^. Intestinal stem cells proliferate in the crypt base and as cells differentiate they move up to the villus, where they can assist in intestinal functions^47,48^. Our IOs are largely composed of intestinal stem cells (immunofluorescence in Fig. 1n) and proliferate quicker than BLOs, demonstrated by the need for passaging IOs and not BLOs (Fig. 1i,1n).

In our comparative analysis of lung and intestinal organoids, we find key differences in mitochondrial organization and behavior. Mitochondrial fragments decreased in size (Fig. 4i) and increased in motility (Fig. 4p) in IOs, but not to the extent that they did in BLOs. Additionally, mitochondria demonstrated more networking in IOs than in BLOs (Fig. 4n,4o). Overall, phenotypes of the mitochondrial network in IOs displayed some similarities with stem cells and some similarities with BLOs. Mitochondrial morphology changed with maturity, where fragments decreased in length with increased cell maturation (Fig. 4h,4l). Mitochondrial motility changed with organoid type, where the least mature lung cells were the most similar to intestinal cells (rows of color in Fig. 4s,4w) and most intestinal ages demonstrated phenotypes more similar to stem cells and DE cells (Fig. 4v, dark rectangle in lower left of Fig. 4s,4w). This data supports a model where mitochondrial phenotype, specifically fragment size and motility, is linked to a cell’s proliferation rate.

Our third finding is that the mitochondrial network encodes enough information about cell state and cell type that it can be used for accurate predictions. We achieved an 83.7% accuracy in predicting organoid type and developmental age using a combination of morphological and temporal features (Fig. 6h,6m). This suggests that morphological and temporal characteristics are strong indicators of cellular identity and maturity. Effectively, our results show that the 4D mitochondrial network is a non-invasive biomarker that reports on cell differentiation status, cell state, and cell function. Importantly, our results show that including both structural and temporal mitochondrial features improves classification accuracy, emphasizing the need for 4D cell biology in developmental studies.

Despite the advances and results that we could obtain using 4D single cell mitochondria quantification in live organoid tissues, our study has limitations. While our segmentation pipeline is robust, further automation and deep learning integration could enhance efficiency and accuracy. Future work should also explore the functional implications of mitochondrial changes, such as their impact on ATP production, reactive oxygen species generation, and metabolic adaptation. The addition of dyes or other fluorescent reporters could provide a reliable readout for these functional characteristics.

In conclusion, our study uses a comprehensive framework for 4D single-cell mitochondrial analysis in human organoids to create the first comprehensive map of how mitochondrial phenotypes are linked to cell state, cell type, and cell fate through development from hiPSCs to mature lung and intestinal organoids. By quantifying mitochondrial dynamics across development, we reveal key similarities and differences between lung and intestinal organoids and demonstrate the potential for mitochondrial metrics to serve as biomarkers of cellular state. These findings lay the groundwork for future investigations into mitochondrial and other organelle function in organ development, disease modeling, and drug discovery using human organoid model systems.

## Methods

### Cell and organoid culture

#### Stem cell culture

hiPSCs endogenously tagged with Tom20-mEGFP were purchased from the Allen Institute for Cell Science through the UCB Cell Culture Facility (AICS-0011 cl.27). The cell membrane was tagged in our lab using the CRISPR-Cas9 system with CAAX-TagRFP (Addgene plasmid #107580a) to create our “CT20” cell line. Stem cells were cultured using mTeSR Plus culture media (STEMCELL Technologies #100-0276) with 1% Pen/Strep and routinely passaged with Accutase and ROCK inhibitor for single cell passaging or ReLeSR for clump passaging. Plates were coated with Matrigel. In preparation for organoid culture, stem cells were clump passaged according to protocols listed below. Cells were routinely tested for mycoplasma and karyotyped.

#### Branching lung organoid culture

BLOs were cultured using STEMCELL Technologies’ STEMdiff Branching Lung Organoid kit (#100-0195). Briefly, CT20s were clump passaged and grown to 30-60% confluency in a 24-well plate for definitive endoderm (DE) differentiation (Stage 1). The definitive endoderm was formed as a dense monolayer of cells and three-dimensional structures began to form (day 0-3). Anterior foregut endoderm buds floated into the media from the monolayer (day 3-6). These were removed and further differentiated into lung bud organoids in suspension culture where they developed more defined branching structures of epithelial cells (day 6-14). Lung bud organoids were differentiated into BLOs in a Matrigel (Corning, 356231) sandwich within a Transwell insert, which allowed for media exchange above and below the BLOs (day 14-42+). They were maintained for 130+ days and expressed lung progenitor marker NKX2-1.

#### Intestinal organoid culture

Intestinal organoids were cultured using STEMCELL Technologies’ STEMdiff Intestinal Organoid kit (#05140). Briefly, CT20s were clump passaged and grown to 85-90% confluency in a 24-well plate for definitive endoderm (DE) differentiation where they began to show three-dimensional structures (Stage 1). During DE differentiation, the cells appeared bigger and more compact, starting in the middle of the dense colonies. After three days of DE culture, mid-/hindgut (MH) differentiation was induced (Stage 2). During MH differentiation, buds came off of the thick monolayer and were suspended in the culture media. These buds were collected and seeded in a Matrigel dome (Stage 3). The length of time buds were exposed to MH media determined what part of the intestine they recapitulated. The buds embedded on earlier days mimicked tissue in the upper intestinal tract (duodenum) and the buds embedded on later days mimicked tissue in the lower intestinal tract (ileum). For uniformity, all organoids used in these experiments were embedded on the same day. Due to the highly proliferative nature of intestinal cells, the organoids quickly expanded to occupy the provided space. After 7-10 days of growth, IOs were passaged. For passaging, cold DMEM was sprayed directly onto the dome to dislodge it. Using a 1mL pipette, IOs were broken up and distributed across 2-5 cold tubes, depending on organoid density. Supernatant in these tubes was disposed of and cold, fresh DMEM was sprayed into each tube. Organoids were spun down and supernatant was disposed of. IO fragments were resuspended in 40µL Matrigel and placed into a 24-well plate in a dome. Growth media was added after the domes had solidified, approximately after 15-30 minutes at 37ºC in the incubator. Organoids were passaged 5+ times over the course of 90+ days in culture and expressed epithelial adhesion marker EPCAM and intestinal epithelial marker CDX2.

#### Lattice light-sheet imaging

A custom built lattice light sheet microscope designed by Eric Betzig’s Lab at HHMI Janelia/UC Berkeley was used to image samples. Key modifications of this work are the use of 0.6 NA excitation objective lens (Thorlabs, TL20X-MPL), a 1.0 NA detection objective lens (Zeiss W Plan-Apochromat 20x/1.0, model #421452-9800), and a Hamamatsu Photonics Orca Fusion BT sCMOS Camera for image acquisition. 488nm and 560nm lasers were used to excite GFP and RFP. A Multiple Bessel Beam Light Sheet Pattern with NA Max 0.4, NA Min 0.35 was used for all samples. A 50msec exposure time was used per 2D scan and samples were scanned every 300nm continuously with a piezo-actuated stage.

All samples were imaged on #1.5 22mm x 22mm square coverslips. Prior to seeding samples, coverslips were sterilized with 100% EtOH and UV light. After sterilization, they were rinsed with 1X DPBS and left to return hydrophilicity for at least 2 hours in the fume hood.

Samples were imaged in phenol red-free DMEM/F12 (Gibco, 11320033) supplemented with 2% FBS (Genesee Scientific 25-514H). The imaging chamber was filled with imaging media and kept at 37ºC and 5% CO2.

#### Monolayer samples

Sterilized coverslips were coated with Matrigel. CT20s were clump passaged onto coverslips and maintained in a 6-well plate until ready for imaging. Differentiation was performed in this 6-well plate according to STEMCELL protocols, with increased media volumes. Day 0 (stem cells) samples were imaged at confluency used to induce DE differentiation. Imaging targets were the middle of colonies with consistent cell confluency. Day 1 (DE lung, intestine) samples were imaged approximately 24 hours post induction. Day 2 (DE lung, intestine) samples were imaged approximately 24 hours post media change. Imaging targets for days 1 and 2 were the middle of colonies where cells were visibly larger.

#### Branching lung organoids

The Matrigel sandwich in the Transwell plate was dissociated using cold DMEM and a cut 1mL pipette tip. Organoids were isolated from each well and placed into separate 1.7mL tubes. Repeated pipetting with cold DMEM and a cut tip was performed to dissociate Matrigel left on the organoid while ensuring organoid tissue stayed intact. Any media left in the tube was removed. Using a cut 200µL tip, the organoid was resuspended in 30µL cold Matrigel and seeded into a dome in the middle of a sterilized coverslip. Matrigel was left for approximately 20 minutes at 37ºC in the incubator until visibly set. Normal culture media was added to cover the dome until ready for imaging, typically overnight. Approximately 30 minutes before imaging, culture media was replaced with phenol red free imaging media to extract phenol red from the Matrigel.

#### Intestinal organoids

Intestinal organoids were passaged as described in the protocol with some modifications. Organoids were passaged onto sterilized coverslips in 30µL Matrigel domes. Normal culture media was added to cover the dome until ready for imaging, typically several days. Approximately 30 minutes before imaging, culture media was replaced with phenol red free imaging media to extract phenol red from the Matrigel.

### Immunostaining

#### Monolayers

Immediately after imaging, samples were rinsed with PBS and fixed with 4% PFA for 20 minutes at room temperature. Samples were then rinsed 3 times with PBS and stored at 4°C in 15% sucrose solution.

For staining, we washed the cells 3 times, 10 minutes each with 1X DPBS at room temperature. We added the permeabilization and blocking buffer and incubated the sample for 1 hour at room temperature. We added the primary antibody cocktail and incubated the sample overnight at 4ºC. The next day, we washed the cells 3 times, 5 minutes each with 1X DPBS at room temperature. We added the secondary antibody cocktail and incubated the sample covered, at room temperature for 2 hours. Finally, we washed the cells again 3 times, 10 minutes each with 1X DPBS at room temperature. We visualized the staining with the ECHO Revolve.

Permeabilization/Blocking (PB) buffer (in 1X DPBS)

- 0.3% Triton X-100
- 2.5% Donkey serum
- 1% BSA

Primary antibody cocktail (in PB buffer):

- 1:400 FoxA2/HNF3B, Rabbit (CellSignaling, #8186)
- 1:200 SOX2, Mouse (Invitrogen, MA1-014)

Secondary antibody cocktail (in PB buffer):

- 1:500 Donkey anti-rabbit, 647nm (Invitrogen, A32795TR)
- 1:500 Donkey anti-mouse, 560nm (Invitrogen, A10037)
- 2 drops/mL NucBlue Fixed Cell Stain ReadyProbes (Invitrogen, R37606)

#### Organoids

Immediately after imaging, samples were rinsed with PBS and fixed with 4% PFA for 60 minutes at room temperature. Samples were then rinsed 3 times with PBS and stored at 4°C in 15% sucrose solution. They were kept on the coverglass on which they were imaged.

Since our samples are grown from our endogenously tagged CT20 cell line, we expose our fixed samples to a 470nm light source to photobleach the endogenous tags and ensure the signal we get from staining originated from our proteins of interest. UV light was shown on the samples for 3-6 hours at room temperature, until minimal fluorescent signal was visible in our ECHO Revolve microscope.

#### BLOs

BLOs were transferred to 30% sucrose solution until the organoids were saturated, a few hours. Organoids were embedded in OCT and placed on dry ice. Organoids were cryosectioned and individual slices were mounted on slides for staining.

Primary antibody cocktail:

- 1:50 NKX2-1, Mouse (Abnova, H00007080-M01)

Secondary antibody cocktail (in PB buffer):

- 1:500 Donkey anti-mouse, 560nm (Invitrogen, A10037)
- 2 drops/mL NucBlue Fixed Cell Stain ReadyProbes (Invitrogen, R37606)

#### IOs

Protocol was adapted from a published protocol^49^. Briefly, stored organoids were washed 3 times for 5 minutes with 1X DPBS. They were washed with EOWB for 15 minutes at room temperature. Liquid was removed and replaced with EOBB. Sample incubated for 1 hour at room temperature. Liquid was removed and replaced with the primary antibody cocktail. Sample incubated overnight at +4ºC. Cocktail was removed and replaced with EOWB. Three EOWB washes were performed at room temperature, with 2 hours for each wash. After the final wash, liquid was replaced with the secondary antibody cocktail. Sample was covered and incubated overnight at +4ºC. Cocktail was removed and replace with EOWB. Three EOWB washes were performed at room temperature, with 2 hours for each wash. We visualized the staining weight the ECHO Revolve.

EOBB (in sterile MilliQ water):

- 10% DPBS 10X
- 0.3% Triton X-100
- 2.5% Donkey serum
- 1% Gelatin
- 1% BSA

EOWB (in sterile MilliQ water):

- 10% DPBS 10X
- 0.3% Triton X-100

Primary antibody cocktail (in EOWB):

- 1:100 CDX2
- 1:100 EPCAM

Secondary antibody cocktail (in EOWB):

- 1:500 Donkey anti-rabbit, 647nm (Invitrogen, A32795TR)
- 1:500 Donkey anti-mouse, 560nm (Invitrogen, A10037)
- 2 drops/mL NucBlue Fixed Cell Stain ReadyProbes (Invitrogen, R37606)

#### Data Preprocessing

LLSM data was preprocessed using methods described in Wang, Hakozaki, et al (2024). Preprocessed data was 974×1536×270px (∼108×171×30µm, xyz) and included 2 color channels (mitochondria in 488nm, cell membrane in 560nm).

#### Cell segmentation and tracking

Preprocessed LLSM data was segmented using our 4D cell segmentation pipeline. The top and bottom 20 z-stacks from the datasets were removed because the data is typically too blurry to obtain reliable segmentations. For every 3D frame (xyz), Cellpose was used to segment cells on each 2D slice (xy) for the membrane channel. The 2D cell masks were then stacked into 3D cell masks using DBSCAN clustering on the centroids. To smooth the edge, a gaussian kernel with sigma 7 was applied on the 3D segmented masks.

To achieve temporal cell tracking, we computed Euclidean distances every 10 z-slices between each centroid label *l* in frame *t* and all centroid labels in frame *t+1*. For each comparison, we computed the mean distance, and the centroid in frame *t+1* with the lowest average distance was assigned to centroid *l*. This process was repeated for all consecutive frames to generate the final 4D cell segmentations. To ensure the quality of the temporal segmentation, poorly segmented cells were manually removed. The 4D cell masks obtained from the membrane segmentation pipeline were then applied to the mitochondrial data to extract single-cell mitochondrial movies. Mitochondrial signal was normalized and converted to 8-bit. We used MitoGraph with global thresholding to segment the mitochondrial surface and obtain skeletons. The binary mitochondrial segmentation output was used to provide numbers of pixels belonging to cell membrane, mitochondria, and cytoplasm and to provide average distances from mitochondria to centroid and cell membrane. The mitochondrial skeleton output was used by MitoTNT to quantify mitochondrial morphology and dynamics.

## Acknowledgements

This study was supported through an NIH Director’s New Innovator Award to JS and through an NIH NIBIB Award to GM under award number T32EB009380.

The authors thank the members of the Schöneberg Lab for helpful discussions. The authors thank Peter Fenton and Loren Looger for assistance in immunostaining branching lung organoids. Organoid development icons were created by Andre Modolo in the Schöneberg Lab.

## Contributions

Gillian McMahon (GM) performed hiPSC differentiation into organoids, prepared samples for LLSM imaging, conceptualized cell tracking algorithm, and conducted data analysis. Dhruv Agarwal (DA) assembled the 4D cell segmentation pipeline and wrote the tracking algorithm. Mehul Arora (MA) wrote the 3D cell segmentation algorithm. Johannes Schöneberg (JS) wrote the cell smoothing algorithm. Zichen Wang (ZW) wrote mitochondria data compilation code. Hiroyuki Hakozaki (HH) conducted LLSM imaging. GM, DA, and JS wrote the manuscript. JS was responsible for conceptualization, funding, and administration.

## Ethics declarations

The authors declare no competing interests.

## References

1. Friedman, J. R. & Nunnari, J. Mitochondrial form and function. Nature 505, 335–343 (2014).

2. Lane, N. & Martin, W. The energetics of genome complexity. Nature 467, 929–934 (2010).

3. Osellame, L. D., Blacker, T. S. & Duchen, M. R. Cellular and molecular mechanisms of mitochondrial function. Best Pract. Res. Clin. Endocrinol. Metab. 26, 711–723 (2012).

4. Mills, E. L., Kelly, B. & O’Neill, L. A. J. Mitochondria are the powerhouses of immunity. Nat. Immunol. 18, 488–498 (2017).

5. Casanova, A., Wevers, A., Navarro-Ledesma, S. & Pruimboom, L. Mitochondria: It is all about energy. Front. Physiol. 14, 1114231 (2023).

6. Liu, Y. J., Sulc, J. & Auwerx, J. Mitochondrial genetics, signalling and stress responses. Nat. Cell Biol. 27, 393–407 (2025).

7. Aghapour, M. et al. Mitochondria: at the crossroads of regulating lung epithelial cell function in chronic obstructive pulmonary disease. Am. J. Physiol. - Lung Cell. Mol. Physiol. 318, L149–L164 (2020).

8. Seo, B. J., Yoon, S. H. & Do, J. T. Mitochondrial Dynamics in Stem Cells and Differentiation. Int. J. Mol. Sci. 19, 3893 (2018).

9. Prowse, A. B. J. et al. Analysis of Mitochondrial Function and Localisation during Human Embryonic Stem Cell Differentiation In Vitro. PLOS ONE 7, e52214 (2012).

10. Ludikhuize, M. C. et al. Mitochondria Define Intestinal Stem Cell Differentiation Downstream of a FOXO/Notch Axis. Cell Metab. 32, 889-900.e7 (2020).

11. Slack, J. M. Intestine in the lung. J. Biol. 3, 10 (2004).

12. Shahbazi, M. N. Mechanisms of human embryo development: from cell fate to tissue shape and back. Dev. Camb. Engl. 147, dev190629 (2020).

13. Muhr, J., Arbor, T. C. & Ackerman, K. M. Embryology, Gastrulation. in StatPearls (StatPearls Publishing, Treasure Island (FL), 2025).

14. Ikonomou, L. & Kotton, D. N. Derivation of Endodermal Progenitors From Pluripotent Stem Cells. J. Cell. Physiol. 230, 246–258 (2015).

15. Agathocleous, M. et al. Metabolic differentiation in retinal cells. Nat. Cell Biol. 14, 859–864 (2012).

16. Collins, T. J., Berridge, M. J., Lipp, P. & Bootman, M. D. Mitochondria are morphologically and functionally heterogeneous within cells. EMBO J. 21, 1616–1627 (2002).

17. Bao, L.-L. et al. Epithelial OPA1 links mitochondrial fusion to inflammatory bowel disease. Sci. Transl. Med. 17, eadn8699 (2025).

18. Tweedy, J. et al. 3D Reconstruction of the Mitochondrial Network within the Neuronal Soma from SBF-SEM Volume Data. Methods Mol. Biol. Clifton NJ 2831, 145–177 (2024).

19. Wang, Z. et al. MitoTNT: Mitochondrial Temporal Network Tracking for 4D live-cell fluorescence microscopy data. PLOS Comput. Biol. 19, e1011060 (2023).

20. Lefebvre, A. E. Y. T., Ma, D., Kessenbrock, K., Lawson, D. A. & Digman, M. A. Automated segmentation and tracking of mitochondria in live-cell time-lapse images. Nat. Methods 18, 1091–1102 (2021).

21. Lefebvre, A. E. Y. T. et al. Nellie: automated organelle segmentation, tracking and hierarchical feature extraction in 2D/3D live-cell microscopy. Nat. Methods 1–13 (2025) doi:10.1038/s41592-025-02612-7.

22. Yang, S. et al. Organoids: The current status and biomedical applications. MedComm 4, e274 (2023).

23. Gu, Y. et al. Organoid assessment technologies. Clin. Transl. Med. 13, e1499 (2023).

24. Lukonin, I., Zinner, M. & Liberali, P. Organoids in image-based phenotypic chemical screens. Exp. Mol. Med. 53, 1495–1502 (2021).

25. Fei, K., Zhang, J., Yuan, J. & Xiao, P. Present Application and Perspectives of Organoid Imaging Technology. Bioengineering 9, 121 (2022).

26. Chen, B.-C. et al. Lattice Light Sheet Microscopy: Imaging Molecules to Embryos at High Spatiotemporal Resolution. Science 346, 1257998 (2014).

27. Liu, T.-L. et al. Observing the cell in its native state: Imaging subcellular dynamics in multicellular organisms. Science 360, eaaq1392 (2018).

28. Schöneberg, J. et al. 4D cell biology: big data image analytics and lattice light-sheet imaging reveal dynamics of clathrin-mediated endocytosis in stem cell–derived intestinal organoids. Mol. Biol. Cell 29, 2959–2968 (2018).

29. Wang, Z., Hakozaki, H., McMahon, G., Medina-Carbonero, M. & Schöneberg, J. LiveLattice: Real-time visualisation of tilted light-sheet microscopy data using a memory-efficient transformation algorithm. J. Microsc. 297, 123–134 (2025).

30. Stringer, C., Wang, T., Michaelos, M. & Pachitariu, M. Cellpose: a generalist algorithm for cellular segmentation. Nat. Methods 18, 100–106 (2021).

31. Ester, M., Kriegel, H.-P., Sander, J. & Xu, X. A density-based algorithm for discovering clusters in large spatial databases with noise. in Proceedings of the Second International Conference on Knowledge Discovery and Data Mining 226–231 (AAAI Press, Portland, Oregon, 1996).

32. Viana, M. P., Lim, S. & Rafelski, S. M. Chapter 6 - Quantifying mitochondrial content in living cells. in Methods in Cell Biology (ed. Paluch, E. K.) vol. 125 77–93 (Academic Press, 2015).

33. McGrath, P. S. & Wells, J. M. SnapShot: GI Tract Development. Cell 161, 176-176.e1 (2015).

34. Baulies, A., Angelis, N. & Li, V. S. W. Hallmarks of intestinal stem cells. Development 147, dev182675 (2020).

35. Fu, W., Liu, Y. & Yin, H. Mitochondrial Dynamics: Biogenesis, Fission, Fusion, and Mitophagy in the Regulation of Stem Cell Behaviors. Stem Cells Int. 2019, 9757201 (2019).

36. Chen, H. & Chan, D. C. Mitochondrial Dynamics in Regulating the Unique Phenotypes of Cancer and Stem Cells. Cell Metab. 26, 39–48 (2017).

37. Cho, Y. M. et al. Dynamic changes in mitochondrial biogenesis and antioxidant enzymes during the spontaneous differentiation of human embryonic stem cells. Biochem. Biophys. Res. Commun. 348, 1472–1478 (2006).

38. Chen, W., Zhao, H. & Li, Y. Mitochondrial dynamics in health and disease: mechanisms and potential targets. Signal Transduct. Target. Ther. 8, 1–25 (2023).

39. Lounas, A. et al. A 3D analysis revealed complexe mitochondria morphologies in porcine cumulus cells. Sci. Rep. 12, 15403 (2022).

40. Glancy, B. et al. Mitochondrial reticulum for cellular energy distribution in muscle. Nature 523, 617–620 (2015).

41. Dahl, R. et al. Three-dimensional reconstruction of the human skeletal muscle mitochondrial network as a tool to assess mitochondrial content and structural organization. Acta Physiol. 213, 145–155 (2015).

42. Zhao, J. et al. Mitochondrial dynamics regulates migration and invasion of breast cancer cells. Oncogene 32, 4814–4824 (2013).

43. Cunniff, B., McKenzie, A. J., Heintz, N. H. & Howe, A. K. AMPK activity regulates trafficking of mitochondria to the leading edge during cell migration and matrix invasion. Mol. Biol. Cell 27, 2662–2674 (2016).

44. Paupe, V. & Prudent, J. New insights into the role of mitochondrial calcium homeostasis in cell migration. Biochem. Biophys. Res. Commun. 500, 75–86 (2018).

45. Hollenbeck, P. J. & Saxton, W. M. The axonal transport of mitochondria. J. Cell Sci. 118, 5411–5419 (2005).

46. Kaunitz, J. D. & Akiba, Y. Control of Intestinal Epithelial Proliferation and Differentiation: The Microbiome, Enteroendocrine L Cells, Telocytes, Enteric Nerves, and GLP, Too. Dig. Dis. Sci. 64, 2709–2716 (2019).

47. Umar, S. Intestinal Stem Cells. Curr. Gastroenterol. Rep. 12, 340–348 (2010).

48. Choi, J. & Augenlicht, L. H. Intestinal stem cells: guardians of homeostasis in health and aging amid environmental challenges. Exp. Mol. Med. 56, 495–500 (2024).

49. Martinez-Ordoñez, A. et al. Whole-mount staining of mouse colorectal cancer organoids and fibroblast-organoid co-cultures. STAR Protoc. 4, 102243 (2023).

